# Protective and aggressive bacterial subsets and metabolites modify hepatobiliary inflammation and fibrosis in PSC

**DOI:** 10.1101/2021.10.31.466701

**Authors:** Muyiwa Awoniyi, Jeremy Wang, Billy Ngo, Vik Meadows, Jason Tam, Amba Viswanathan, Yunjia Lai, Stephanie A. Montgomery, Morgan Farmer, Martin Kummen, Louise B. Thingholm, Christoph Schramm, Corinna Bang, Andre Franke, Bernd Schnabl, Kun Lu, Jenny PY Ting, Yuri V. Popov, Johannes R. Hov, Heather Francis, R. Balfour Sartor

## Abstract

**Objective:** Conflicting microbiota data exist for primary sclerosing cholangitis (PSC) and experimental models. <u>Goal:</u> Define complex interactions between resident microbes and their association in PSC patients by studying antibiotic-treated specific pathogen-free (SPF) and germ-free (GF) multi-drug-resistant 2 deficient (*mdr2*^*-/-*^) mice.

**Design:** We measured weights, liver enzymes, RNA expression, histological, immunohistochemical and fibrotic biochemical parameters, fecal 16s rRNA gene profiling, and metabolomic endpoints in gnotobiotic and antibiotic-treated SPF *mdr2*^*-/-*^ mice and targeted metagenomic analysis in PSC patients.

**Results:** GF *mdr2*^*-/-*^ mice had exaggerated hepatic inflammation and fibrosis with 100% mortality by 8 weeks; early SPF autologous stool transplantation rescued liver-related mortality. Broad-spectrum antibiotics and vancomycin alone accelerated disease in weanling SPF *mdr2*^*-/-*^ mice, indicating that vancomycin-sensitive resident microbiota protect against hepatobiliary disease. Vancomycin treatment selectively decreased Lachnospiraceae and short-chain fatty acids (SCFAs) but expanded Enterococcus and Enterobacteriaceae. Antibiotics increased cytolysin-expressing *E. faecalis* and *E. coli* liver translocation; colonization of gnotobiotic *mdr2*^*-/-*^ mice with translocated E. *faecalis* and *E. coli* strains accelerated liver inflammation and mortality. Lachnospiraceae colonization of antibiotic pre-treated *mdr2*^*-/-*^ mice reduced liver fibrosis, inflammation and translocation of pathobionts, while Lachnospiraceae-produced SCFA decreased fibrosis. Fecal *E. faecalis*/ Enterobacteriaceae was positively and Lachnospiraceae was negatively associated with PSC patients’ clinical severity Mayo risk scores.

**Conclusions:** We identified specific functionally protective and detrimental resident bacterial species in *mdr2*^*-/-*^ mice and PSC patients with associated clinical outcomes. These insights may guide personalized targeted therapeutic interventions in PSC patients.

## Introduction

Idiopathic primary sclerosing cholangitis (PSC) is characterized by chronic biliary inflammation, cirrhosis, and variable progression to liver failure^1^. Mechanisms driving the onset and progression of PSC are unclear, contributing to the lack of established therapies to prevent disease progression. Several studies show the prevalence of dysbiotic microbiota in PSC patients with or without IBD characterized by decreased microbial diversity and over-represented intestinal pathobionts like Enterococcus *(E. faecalis)*, Enterobacteriaceae *(E. coli, Klebsiella pneumonia*) *and Veillonella*, predominance of biliary *E. faecalis* compared to controls suggesting potential translocation^2,3,4^,^5^ and drive of interleukin (IL)-17-mediated mucosal immunity^2,6^,^7, 5^. In parallel, PSC patients exhibit decreased abundance of presumed protective fecal Lachnospiraceae^8,4^, with bile acid (BA) metabolic activities and anti-colitogenic properties^9^ but unclear functional hepatobiliary impact. Therapeutic strategies using traditional Lactobacillus and Bifidobacterium probiotic strains^10^ or single donor FMT^11^ are ineffective for short-term treatment in PSC patients, possibly related to lack of colonization of these bacteria in PSC patients^3 4 8^. More targeted hepatoprotective bacterial enrichment strategies are lacking. These studies detail the complexity and strong association of microbiota and PSC but also show the need for further investigation of potential pathogenic and protective functional roles of resident gut microbiota in advancing understanding of PSC pathogenesis.

Preclinical studies document the protective role of unfractionated gut microbiota in PSC. Germ-free (GF) C56BL/6 wild-type (WT) mice with chemically induced liver fibrosis models (CCl_4_ and TAA)^12^ or GF FVB/N multi-drug resistance 2 deficient (*mdr2*^*-/-*^) mice^13^, a widely studied model of spontaneously developing PSC, exhibit accelerated periductal fibrosis and progressive cholestasis in the absence of microbes compared to conventionally raised controls. Accelerated biliary inflammation in *mdr2*^*-/-*^ mice is presumed due to toxicity of non-micellar BA in the bile duct^14^. Dysbiosis in PSC patients may result in altered BA pools that potentially contribute to the PSC pathophysiology^3,15^. More recently, increased microbial bile salt hydrolase (BSH) activity was found in bile of PSC patients^3^. *E. faecalis*, abundant in the stool of PSC patients^4,3^, have high BSH activity^16,17^. These observations suggest a potential pathogenic role for bacterial BSH activity in PSC. Functional microbial or metabolic studies to further delineate mechanistic differences amongst intestinal bacteria that selectively influence hepatobiliary inflammation and fibrosis in PSC are lacking.

To address the uncertain functional roles of resident bacteria in cholestatic liver disease, we utilized gnotobiotic and antibiotic approaches in specific pathogen free (SPF) *mdr2*^*-/-*^ mice. We independently identified functionally protective (Lachnospiraceae) and pathogenic (*E. faecalis* and *E. coli*) resident bacteria and defined *in vivo* mechanisms through complementation studies. Conserved abundance of these bacteria were found in metagenomic analysis of PSC patient feces; with unique evidence of a moderate association with clinical outcomes in PSC patients denoting conserved microbial roles in experimental and clinical disease. These insights could improve targeted donor selection on future PSC-FMT trials and drive selective fecal enrichment/depletion approaches to increase efficacy of future personalized microbiotal therapies.

## Methods

### Ethics Statement/animal husbandry

We conducted mouse studies under the NIH guide for Care and Use of Laboratory Animals, approved and overseen by the UNC-Chapel Hill Institutional Animal Care and Use Committee (Protocol ID 18-266). The Popov lab generated congenic C57Bl/6 SPF *mdr2(abcb4*)^*-/-*^ mice^18^. *Mdr2*^*-/-*^ and WT control C57BL/6 mice were housed in microisolator cages with no more than 5 mice/cage in SPF vivaria or GF Trexlar isolators. The UNC National Gnotobiotic Rodent Resource Center (P40OD010995) derived GF *mdr2*^*-/-*^ mice by Caesarian section. We bred GF *mdr2*^*+/-*^ heterozygote mice because of early mortality of GF *mdr2*^*-/-*^ mice. Age and sex-matched mice had unrestricted access to autoclaved water and 5v0F Purina LabDiet chow with a 12-hr light/dark cycle.

### Antibiotics Treatment

Non-fasted 3-4wk old male and female mice received either combined or single antibiotics (0.5mg/ml vancomycin (Hospira), 1mg/ml neomycin (Medisca), and 50 mg/kg metronidazole (G.D. Searle) in drinking water ad libitum, consuming 6-7 milliliters/mouse/day^19,20^. The antibiotic mixture was diluted in deionized H_2_O, sterilized through a 0.2 µm filter and replaced twice weekly for 7-14 days until harvest.

### Assessing of Bacterial Liver Translocation

Left lobe (40mg) was sterilely homogenized in 200μl of PBS. 100μl were plated and serially diluted homogenously and spread onto UTI ChromoSelect Agar (Millipore/Sigma 16636) after initially plating on non-selective BHI plates, incubated overnight (37°C), and colonies were counted. Red-pink and blue colonies representing *E. coli* and *Enterococcus spp* (β-D-galactosidase and ß-glucosidase enzyme activity, respectively), were counted and multiplied by the dilution factor. Sanger sequencing verified colonies as *E. faecalis* or *E. coli*.

### Primary Sclerosing Cholangitis Patient Fecal analysis

Metagenomic analysis of fecal samples collected from 136 patients with PSC and 158 healthy controls from multi-national (Norwegian and German) repositories was performed by Dr. Hov’s group. Antibiotic (Abx) exposure in PSC patients was defined within 6-months of the study (+Abx = 58, no Abx = 24 patients). A detailed patient description is provided in previous study ^4^.

### Statistical Analysis

Murine data shown as mean ± SEM, with “n” representing number of mice indicated in figure legends. One dot or lane represents one mouse. Linear fitting and normalization were performed in GraphPad Prism. Unless specified in the figure legends, statistical significance between two groups was determined by unpaired, two-tailed Student’s t-test; significance between >2 groups were determined using one-way ANOVA with Fisher’s LSD test using GraphPad Prism default settings. We compared CFU numbers by the Poisson generalized linear model, estimated with a Markov Chain Monte Carlo method robust for small sample sizes. Error bar represents mean ± SEM with *p < 0.05, **p < 0.01, ***p < 0.001, ****p < 0.0001 and n.s. no significance. Further details of histological staining, IHC staining and analysis, bacterial or metabolite inoculation and cultivation, biochemical, liver enzyme, metabolomic and 16s rRNA sequencing/analysis are presented in the Supplementary Material.

## Results

### Absent microbiota potentiates hepatobiliary inflammation, fibrosis and mortality in *mdr2*^*-/-*^ mice and is rescued by early autologous stool transplant

To evaluate resident microbiota’s role in this experimental cholestatic murine model, we generated and characterized GF C57BL/6 *mdr2*^*-/-*^ mice compared to littermate C57BL/6 WT mice. *Mdr2*^*-/-*^ C57BL/6 mice have more aggressive liver injury than FVB/n mice^18^. Relative to SPF conditions, 6-7wk old GF *mdr2*^*-/-*^ mice had decreased weight gain, 2x increased alkaline phosphatase (ALP) and 15x increased total bilirubin, (TB)(Fig. 1A-C). GF *mdr2*^*-/-*^ from weaning up to 5-week-old had grossly normal livers and no histologic inflammation or fibrosis (data not shown), including normal baseline TB (1C). Very rapid onset of serum hyperbilirubinemia occurred beyond age 6 weeks, not appreciated in age-matched SPF *mdr2*^*-/-*^ mice or in GF WT C57BL/6 or older GF *mdr2*^*+/-*^ mice (up to 70 weeks)(1C).

**Figure 1:**
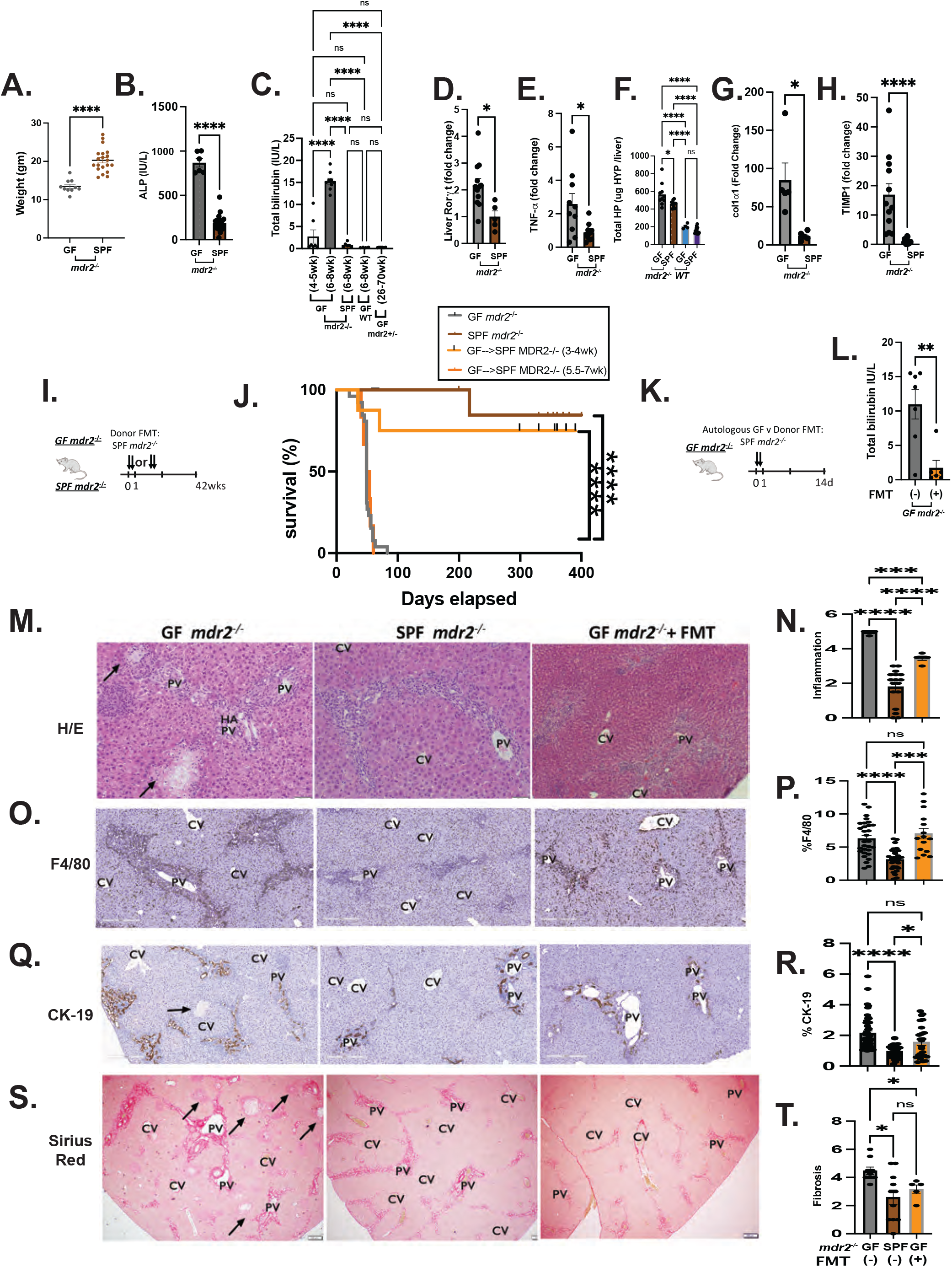
Clinical, biochemical, and liver histological indicators of germ-free *mdr2*^*-/-*^ mice with or without fecal microbiotal transplant compared to control mice. Mouse body weight (A) and serum alkaline phosphatase (AP, B) of 6–8-week germ-free (GF) (N=6-10) and SPF (N=20) *mdr2*^*-/-*^ mice. Longitudinal assessment of biliary injury via serum total bilirubin (TB) in GF *mdr2*^*-/-*^ mice from 4-5 week(N=7) to 6–8-week-old (N=8), compared 6–8-week-old SPF *mdr2*^*-/-*^ and C57/BL6 WT mice and older(26-70 week) old *mdr2* ^*+/-*^ mice (C). Relative expression (fold change) of Rorγt and *tnf-α* (D-E) in livers of 6-8 wk GF (N=5) and SPF (N=5) *mdr2*^*-/-*^ mice. Hepatic collagen deposition expressed as total hepatic hydroxyproline (HYP) expressed in ug HYP/whole liver, calculated by multiplying individual liver weight with relative HYP content (F). Relative expression (fold change) of Collagen I alpha-1 (*col1α1)* and Tissue inhibitor of metalloproteinases (TIMP)-1 (G-H) in livers of 6-8 wk GF (N=5) and SPF (N=5) *mdr2*^*-/-*^ mice. Experimental design of autologous fecal microbiota transplant (FMT) study: Kaplan-Meyer survival curves of pooled Specific pathogen free (SPF) *mdr2*^*-/-*^ donor stool orally gavaged into GF *mdr2*^*-/-*^ C57Bl/6 mice twice in the first 2 days at age 3-4weeks (N=8) or 5.5-7 weeks (N=6) compared to untreated GF (N=23) or SPF (N=14) *mdr2*^*-/-*^ mice monitored for survival for 42 weeks. Experimental design of GF *mdr2*^*-/-*^ given autologous GF (N= 7) or SPF donor (N=6) stool for 14 days (K) monitoring serum TB (L). Representative photomicrographs of 6-8wk old GF *mdr2*^*-/-*^ mice +/-fecal microbiota transplant compared to age-matched SPF *mdr2*^*-/-*^ mice with blinded scoring for liver inflammation (M-N), macrophages (O-P), ductular reaction (Q-R), and fibrosis (S-T) stained by H&E (200X), antibodies to F4/80 (200X) and CK-19 (200X), and Sirius Red (100X), respectively (PV= Portal Vein, CV= Central Vein). Results are expressed as means +/-SEM. Survival data analyzed by Log-rank (Mantel-Cox) test, group or pairwise comparisons performed by ANOVA or Student t-test, respectively. Welch correction was applied to histological scoring analysis. *P < .05, **P < .01, ***P <.001, ****P < .0001

GF *mdr2*^*-/-*^ had increased hepatic RNA expression of the IL-17/IL-22 master regulator, Rorγt, and the inflammatory cytokine *TNF-*α (Fig. 1D,E). Total hepatic hydroxyproline content, a collagen biochemical marker, was markedly increased in GF *mdr2*^*-/-*^ mice, along with RNA expression of fibrinogenic markers, *co1α1* and *timp-1*(Fig. 1F-H). GF and SPF *mdr2*^*-/-*^ mice developed hepatomegaly compared to WT controls (Fig. S1A). No group exhibited intestinal inflammation (not shown). Despite severely elevated serum ALP and TB in GF *mdr2*^*-/-*^ compared to the WT controls, expression of ileal BA transporter, ASBT, FXR or ileal FXR signaling (FGF15) was unchanged (Fig. S1B-E), but expression of hepatic Cyp7a1 was attenuated in GF vs SPF *mdr2*^*-/-*^ mice (S1G).

GF *mdr2*^*-/-*^ mice had markedly decreased survival (median mortality 7.5wks and 100% mortality by 8wks, compared to SPF *mdr2*^*-/-*^ mice with 80% survival past 350d (Fig. 1J). Fecal transplantation of SPF *mdr2*^*-/-*^ feces to 3-4-week-old GF *mdr2*^*-/-*^ mice increased survival (75% at 300 days; Fig. 1J) and improved TB (Fig. 1L) 2 weeks post-therapy. Fecal transplantation later in life (5.5-7wks) provided no protection (Fig. 1J). Although both SPF and GF female *mdr2*^*-/-*^ mice developed more prominent periportal inflammation relative to males, sex-related mortality was not different in GF *mdr2*^*-/-*^ mice. Histologic evaluation of inflammation and ductal reaction examined hematoxylin-eosin (H&E) or anti-F4/80-stained and CK-19 antibody-stained liver sections. GF *mdr2*^*-/-*^ mice displayed heightened periportal inflammation, markedly increased periductular macrophages, lobular necrosis and ductular proliferation (Fig. 1M-R) and more pronounced macrophage (Fig. 1O,P) than neutrophilic (SF1J) infiltration vs SPF *mdr2*^*-/-*^ mice. SPF *mdr2*^*-/-*^ mice displayed more prominent sinusoidal macrophage distribution compared to periductular localization in GF mice, whereas SPF donor FMT-treated GF mdr2^-/-^ had a mixed phenotype. Notable areas of hepatic necrosis (arrows) and granulomatous lesions in GF *mdr2*^*-/-*^ mice stained heavily for F4/80. SPF *mdr2*^*-/-*^ mice developed periductal fibrosis with focal parenchymal extensions, but GF *mdr2*^*-/-*^ mice had doubled histologic fibrosis scores with more extensive periductal fibrosis that extended well into the parenchyma with frequent bridging to portal veins, resembling stage 3/4 patterns of Ishak fibrosis (Fig. 1S,T). FMT attenuated periportal inflammation and fibrosis of GF *mdr2*^*-/-*^ mice (Fig. 1M-N,S-T). No histologic hepatic or ileocolonic inflammation occurred in SPF, GF WT or *mdr2*^*+/-*^ mice. Thus, early life exposure to resident microbiota in GF *mdr2*^*-/-*^ mice have an overall protective and tolerogenic role, decreasing hepatic inflammation, ductal reaction and fibrosis.

### *Mdr2*^*-/-*^ have dysbiotic microbial landscapes with altered metabolic pathways exacerbated by antibiotics

Notable baseline divergence in resident microbial constituency in SPF *mdr2*^*-/-*^ and WT mice by β-diversity analysis (Fig. 2A) with relative decrease in putative protective Clostridial family (Fig. 2B). We developed an antibiotic depletion model in SPF *mdr2*^*-/-*^ mice to validate the protective role of host-microbiota and identify potential beneficial bacterial groups and metabolites. Broad-spectrum antibiotic cocktail (Abx) of vancomycin, neomycin and metronidazole to target resident Gram-positive, Gram-negative, and anaerobic bacteria, respectively^19, 20^ was administered in drinking water ad libitum to 3-4wk old SPF *mdr2*^*-/-*^ mice for 14 days (Fig. 2C). As predicted, Abx similarly depleted microbial abundance in *mdr2*^*-/-*^ and WT control mice by α-diversity and validated fecal qPCR (Fig. 2D-G). Markedly different microbial communities existed in the two hosts following Abx or when assessing effects on mdr2-/- mice alone by *β*- diversity analysis (Fig. 2H-I).

**Figure 2:**
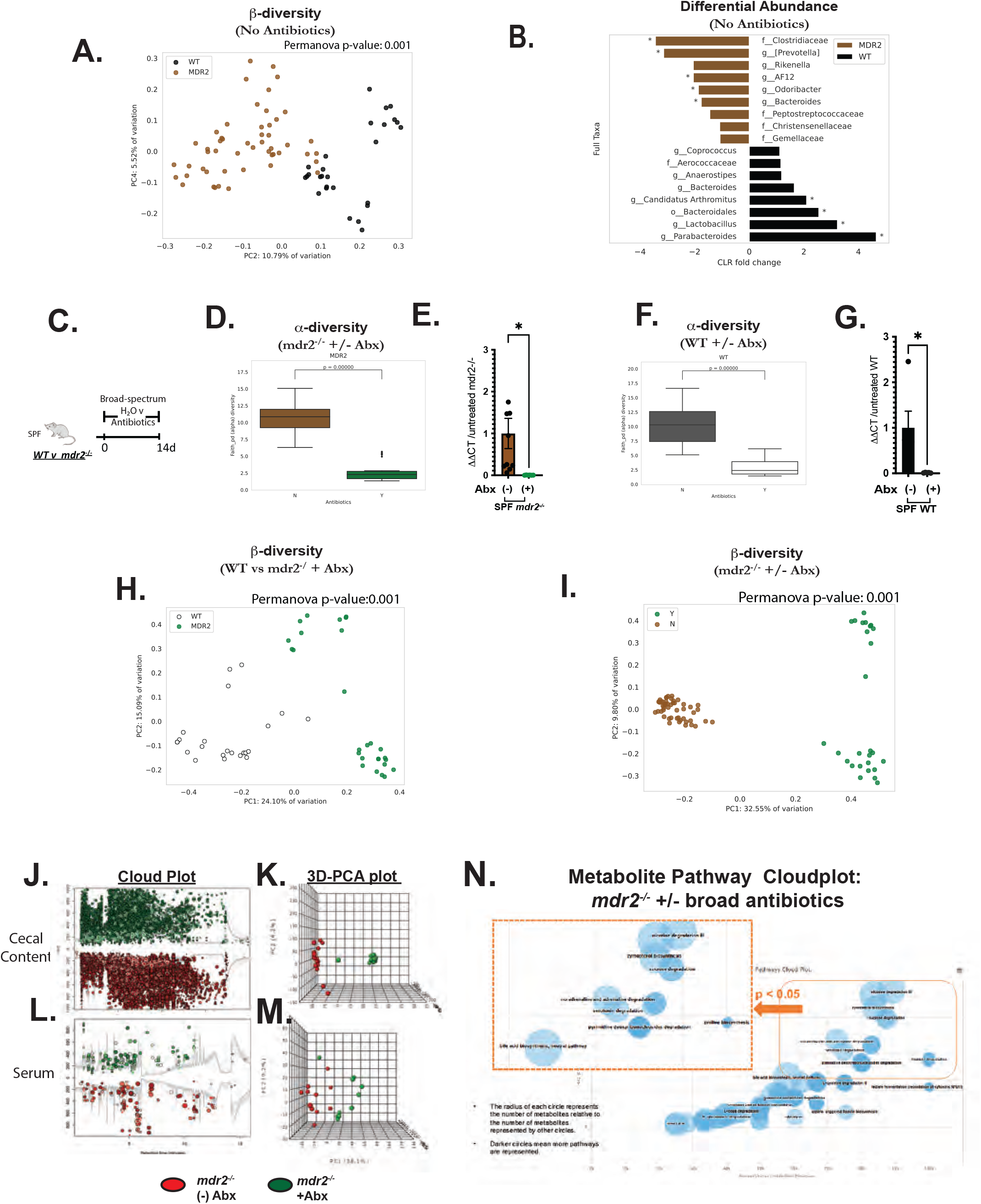
Fecal microbiome and metabolite metabolite profiles of pre/post-antibiotic treated *mdr2*^*-/-*^ and WT mice on C57BL/6 background. Pooled baseline untreated SPF *mdr2*^*-/-*^ and WT C57BL/6 fecal beta diversity (Permanova test) and Linear Discriminant Analysis plot showing differential enrichment of taxa (A-B). Experimental design of broad-spectrum antibiotic (Abx) study, 3-5 week old SPF *mdr2*^*-/-*^ or WT mice exposed ad libitum to broad-spectrum antibiotics (metronidazole, 30mg/ml; vancomycin, 0.5mg/ml; and neomycin 1mg/ml) in drinking H2O or H2O alone (N=10 mice/group) for 14d (pooled from 3 experiments) (C). Effect of 14days of Abx assessing pooled baseline alpha diversity (Faith’s PD) and fecal universal 16s qPCR (expressed in ΔΔCT differences) in *mdr2*^*-/-*^ (D-E) and WT BL/6 (F-G) mice, respectively. PcoA plots assessing beta diversity (Permanova test) antibiotic exposure between genotypes (WT and *mdr2*^*-/-*^ mice) (H) and antibiotic exposure in *mdr2*^*-/-*^ mice (I). AntibiotiCloud Plot of cecal content metabolome with 23730 molecular features with fold changes >1.5 (p<0.01) and the 3D-PCA plot shows the separation of cecal metabolites from *mdr2*^*-/-*^ with and without antibiotics. (J-K) Cloud Plot of serum metabolome with 461 molecular features with fold changes >1.5 (p<0.01and the 3D-PCA plot shows the separation of serum from *mdr2*^*-/-*^ with and without antibiotics. (L-M).)Pathway cloud plot of potential pathway differences with exposure of broad antibiotics in SPF *mdr2*^*-/-*^ mice (N) The radius of each circle represents the number of metabolites relative to the number of metabolites represented by other circles. Darker circles mean more pathways are represented. (*mdr2*^*-/-*^ no abx, N=13, *mdr2*^*-/-*^ with abx, N=25). Group or pairwise comparisons performed by ANOVA or Student t-test, respectively. *P < .05, **P < .01, ***P <.001, ****P < .0001

To identify potential metabolic profiles relevant to protective microbial effects in *mdr2*^*-/-*^ mice, we analyzed the metabolome of serum and cecal contents from antibiotic-treated vs. untreated *mdr2*^*-/-*^ mice by liquid chromatography-high resolution mass spectrometry (Fig. 2J-N). We identified compounds in the most radically changed molecular features (fold change >3, p-value <0.01, Welch’s t-test) to define antibiotic-induced metabolites in cecal contents more accurately^21^. Abx-treatment decreased a spectrum of compounds, spanning BA, alcohols, indoles, oxosteroids, bilirubin, and methyl-branched fatty acid pathways (Fig. 2J-N). We then used these metabolomic profiles to inform our selection of putative protective bacterial candidates that might rescue inflammation/fibrosis driven by Abx depletion of protective microbial subsets.

### Antibiotic depletion of resident microbiota augments restricted weight gain, liver inflammation, biliary epithelial reactivity, and fibrosis in *mdr2*^*-/-*^ mice

Abx depleted the putative protective bacterial families, Clostridiaceae and Lachnospiraceae, by fecal 16s analysis (Fig. 3B). Weight loss in some mice after 14d treatment limited the duration of Abx exposure (Fig. 3C), with increased serum ALP (Fig. 3D) and clinical illness (lethargy, hunching, unkempt fur and peritoneal jaundice), but not robust serum bilirubin elevation (Fig. 3E)^15^. No change in weight, LFTs, or clinical appearance occurred in WT C56BL/6 mice exposed to these Abx (S1K-L). Similar to GF *mdr2*^*-/-*^ mice, Abx-induced inflammation was documented by increased hepatic *Tnf*-α expression, but no change in *Rorγt* (Fig. 3F). Histological evaluation of Abx-treated *mdr2*^*-/-*^ displayed accelerated cholangiopathic injury similar to GF *mdr2*^*-/-*^ mice with expanded portal and peribiliary inflammation with ductular reaction (Fig. 3J-M).

**Figure 3:**
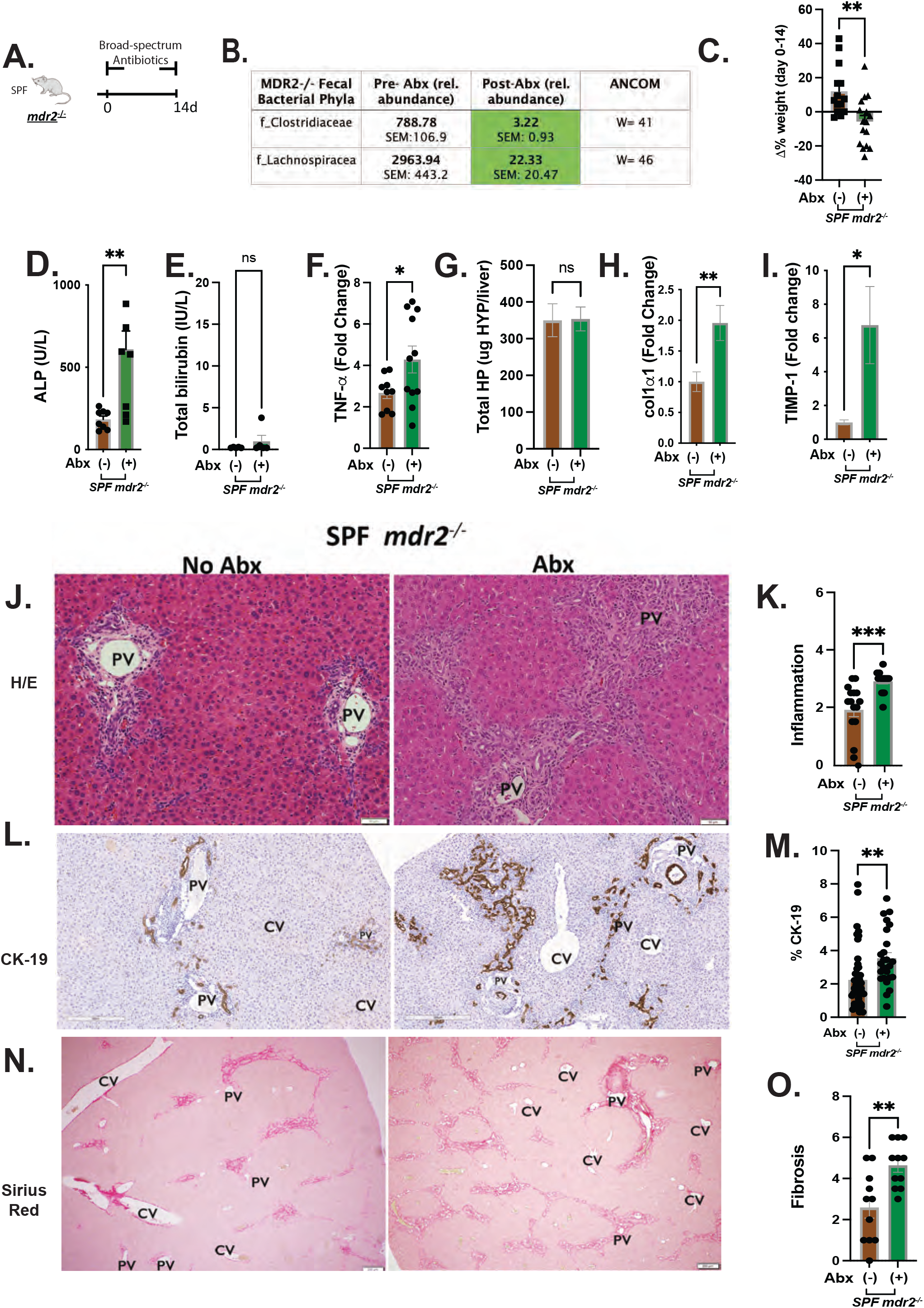
Clinical, biochemical, and liver histological indicators of broad-spectrum antibiotic treated *mdr2*^*-/-*^ mice. Experimental design of broad-spectrum antibiotic (Abx) study, 3–5-week-old SPF *mdr2*^*-/-*^ mice exposed ad libitum to broad-spectrum antibiotics in drinking H2O (N=15) or H2O alone (N=13) for 14d (pooled from 3 experiments) (A). Effect of Abx exposure on various outcomes including relative abundance of selected bacterial phyla expressed as unadjusted raw average OTU relative abundance (B), 14 day body weight change (C), serum ALP, TB (D-E), RNA expression of liver *tnf*-α (F), *col1α1* (H) and *timp*-1 (I) and hepatic collagen deposition expressed as ug HYP/whole liver (G). Representative photomicrographs and blinded composite histologic scoring murine liver stained with H&E (J-K), CK-19 (L-M) and Sirius Red (N-O) of untreated (N=15) vs. Abx-treated SPF *mdr2*^*-/-*^ mice (N=15). Experimental design of treatment of 3-4wk old GF *mdr2*^*-/-*^ mice gavaged twice with pooled *E. faecalis* and *E. coli* hepatic isolates (Ec/Ef) 10^8^ CFU (P), measuring translocated liver bacteria (Q) and blinded liver inflammation score (R) of GF *mdr2*^*-/-*^ mice or water treated controls. Experiment design and Kaplan-Meyer survival curves of 10^8^Ec/Ef vs Lachnospiraceae (Lachno) or H2O controls (S-T) GF *mdr2*^*-/-*^ H2O, N=12; Ec/Ef, N= 5, Lachno, N=6). Heat map representation of cecal content bile acids (U) following Lachno vs H2O control *mdr2*^*-/-*^ accelerated antibiotic model. Results are expressed as means +/-SEM. Pairwise comparisons performed by Student t-test for biochemical and molecular studies; Welch correction was applied to histological scoring analysis. Unadjusted raw average OTU relative abundance and standard errors of bacterial groups against the variables detected with significant effects by ANCOM (analysis of composition of microbes). *P <.05, **P < .01, ***P <.001, ****P < .0001.

Abx-treated *mdr2*^*-/-*^ mice displayed increased lobular collagen deposition often bridging from portal vein to portal vein, compared to untreated SPF *mdr2*^*-/-*^ mice, where fibrosis was limited to periportal areas accompanied by small duct proliferation and focal scarring (Fig. 3N,O). Unchanged hydroxyproline content (Fig. 3G) is likely due to the short study duration, but increased liver RNA expression of *co1α1* and *timp-1* occurred in Abx-treated *mdr2*^*-/-*^ mice vs untreated controls (Fig. 3H-I). Abx did not affect hepatotoxicity or weight gain in WT C57BL/6 mice (S1K-M). This Abx depletion validated our GF mdr2^*-/-*^ findings, confirming that loss of protective bacteria had deleterious effects in this cholestatic model.

### Selective antibiotics have differential effects on *mdr2*^*-/-*^ hepatobiliary disease

To identify microbial population(s) responsible for protecting *mdr2*^*-/-*^ mice, we selectively tested individual Abx components: vancomycin, neomycin, and metronidazole (Fig. 4A, VNM). Vancomycin most significantly decreased fecal bacterial alpha diversity (Fig. 4B) and maintained most divergent bacterial populations with minimal β-diversity overlap with untreated controls (Fig. 4B). Vancomycin and metronidazole decreased the putative protective bacterial family, Clostridaceae, but only vancomycin attenuated Lachnospiraceae (Fig. 4D). Neomycin did not significantly affect either family (Fig. 4D). Vancomycin treatment did not affect bilirubin (4G), similar to Abx, but restricted weight gain and elevated serum ALP, hepatic TNF-α expression, histologic inflammation and ductal reaction (Fig. 4E-F,H-J).

**Figure 4:**
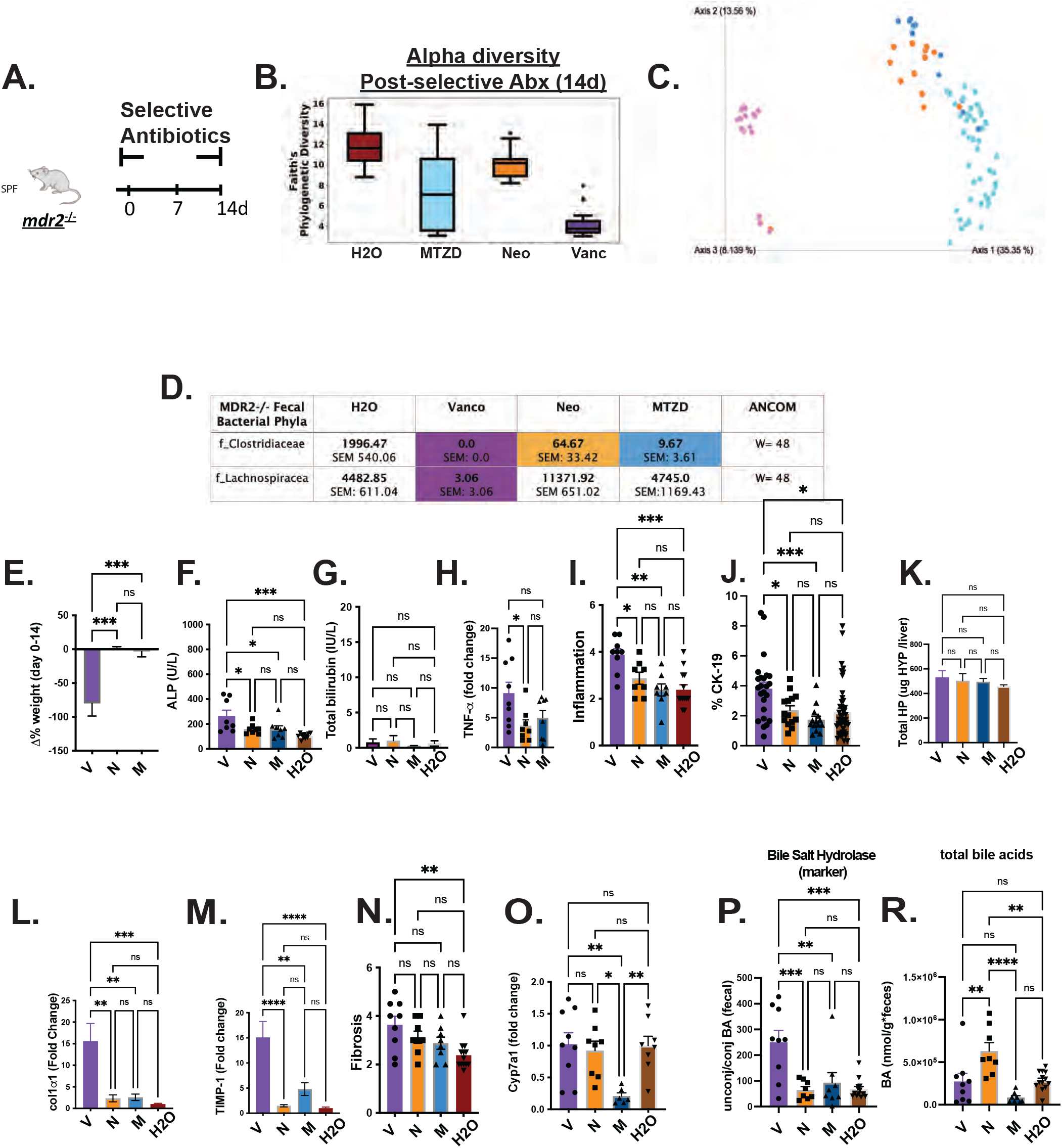
Differential effects of selective antibiotics on *mdr2*^*-/-*^ microbial, clinical, biochemical and histologic outcomes. Experimental design of selective antibiotic treatment (A): SPF *mdr2*^*-/-*^ mice were given individual antibiotics (vancomycin (V, 0.5mg/ml, N=16), neomycin (N, 1mg/ml, N=12), and metronidazole (M, 30mg/ml, N=16)) in autoclaved drinking water ad libitum and control *mdr2*^*-/-*^ mice received autoclaved water alone (N=16) for 14d. Alpha diversity, beta diversity (Permanova test), and differential abundance (Centered Log-Ratio (CLR) transformation from CoDA methods) of Clostridaceae and Lachnospiraceae from 14d fecal samples (B-D). Measured parameters include: 14d weight change (E), serum ALP and TB ()F-G), liver RNA expression of *tnf*-α (H), *col1α1* (J); along with blinded composite histologic scoring murine liver stained with H/E (I), CK-19 (J). Fibrosis readouts include hepatic collagen deposition (K),RNA expression of *timp*-1 (L), *cyp7α1* (M) along with Sirius Red composite staining (N) of selective antibiotic treated *mdr2*^*-/-*^ compared to H_2_O control. (N, Vancomycin=9, Neomycin=8, MTZD=8, H2O=11). *cyp7a1* liver expression (O) as well as pooled cecal content bile acids (BA) differences in surrogate bile salt hydrolase activity indicator based on ratios of total unconjugated/conjugated BA (P) and total BA (R). Group or pairwise comparisons performed by ANOVA or Student t-test, respectively. Welch correction was applied to histological scoring analysis. Unadjusted raw average OTU relative abundance and standard errors of bacterial groups against the variables detected with significant effects by ANCOM (analysis of composition of microbes). *P < .05, **P < .01, ***P <.001, ****P < .0001.

Concurrent with hepatobiliary injury and inflammation, vancomycin exacerbated hepatic fibrogenesis. Relative to water control mice, only vancomycin significantly increased histological liver fibrosis scores (Fig. 4N), expression of hepatic pro-fibrotic *co1α1* and *timp-1* (Fig. 4L-M) but did not change hepatic hydroxyproline (Fig. 4K).

Exploration of VNM on BA metabolism demonstrated isolated metronidazole attenuation of hepatic *cyp7a1* expression SPF *mdr2*^*-*/-^ mice (Fig. 4O). Vancomycin profoundly increased BSH activity, based on the unconjugated/conjugated BA ratio (Fig. 4P), despite not changing total BAs vs control (Fig. 4R). These aggregate data suggest that depletion of vancomycin-sensitive populations, including Clostridiaceae and Lachnospiraceae, results in aggressive hepatobiliary responses.

### Lachnospiraceae (Lachno) administration attenuates phenotypic effects of antibiotic-treated SPF *mdr2*^*-/-*^ mice and inhibits hepatic translocation of *E. faecalis* and *E. coli*

We next tested the ability of putative protective microbial subsets to protect antibiotic-depleted SPF *mdr2*^*-/-*^ mice. To identify triple antibiotic therapy’s shortest duration that accelerated hepatobiliary inflammation/fibrosis but maintained a 2wk experimental window for microbial transfer experiments, we treated 3-4wk old SPF *mdr2*^*-/-*^ mice with Abx for 14-days or 7-days followed by 7d of water (S2A). Interestingly, the shorter 7-day treatment enhanced hepatic inflammation and ALP elevation (S2B-C), without significantly altering fibrosis scores (S2D). We subsequently used accelerated 7d antibiotic treatment to test reconstitution of putative protective bacteria, Clostridiaceae and Lachnospiraceae.

Because Abx depleted known protective fecal Clostridium clusters IV and XIVa^22,23^ in both SPF WT and *mdr2*^*-/-*^ mice (S2E-F), we initially tested a 17 strain consortium of protective human Clostridia^22^. Despite Clostridium strains and their efficacy in multiple colitis models^22^, repletion of antibiotic-pretreated *mdr2*^*-/-*^ mice with these Clostridia did not alter body weight or LFTs (S2G-J). Therefore, we assessed the biological activity of another potentially protective bacterial group depleted by antibiotics, Lachnospiraceae. Serial treatment with a protective colonic 23 strain Lachnospiraceae consortium^24^ (Fig. 5A) restored weight gain and reduced histologic liver inflammation, ductal reaction and fibrosis (Fig. 5B,5E-G), but did not decrease serum biochemistries (Fig. 5C, S2K).

**Figure 5:**
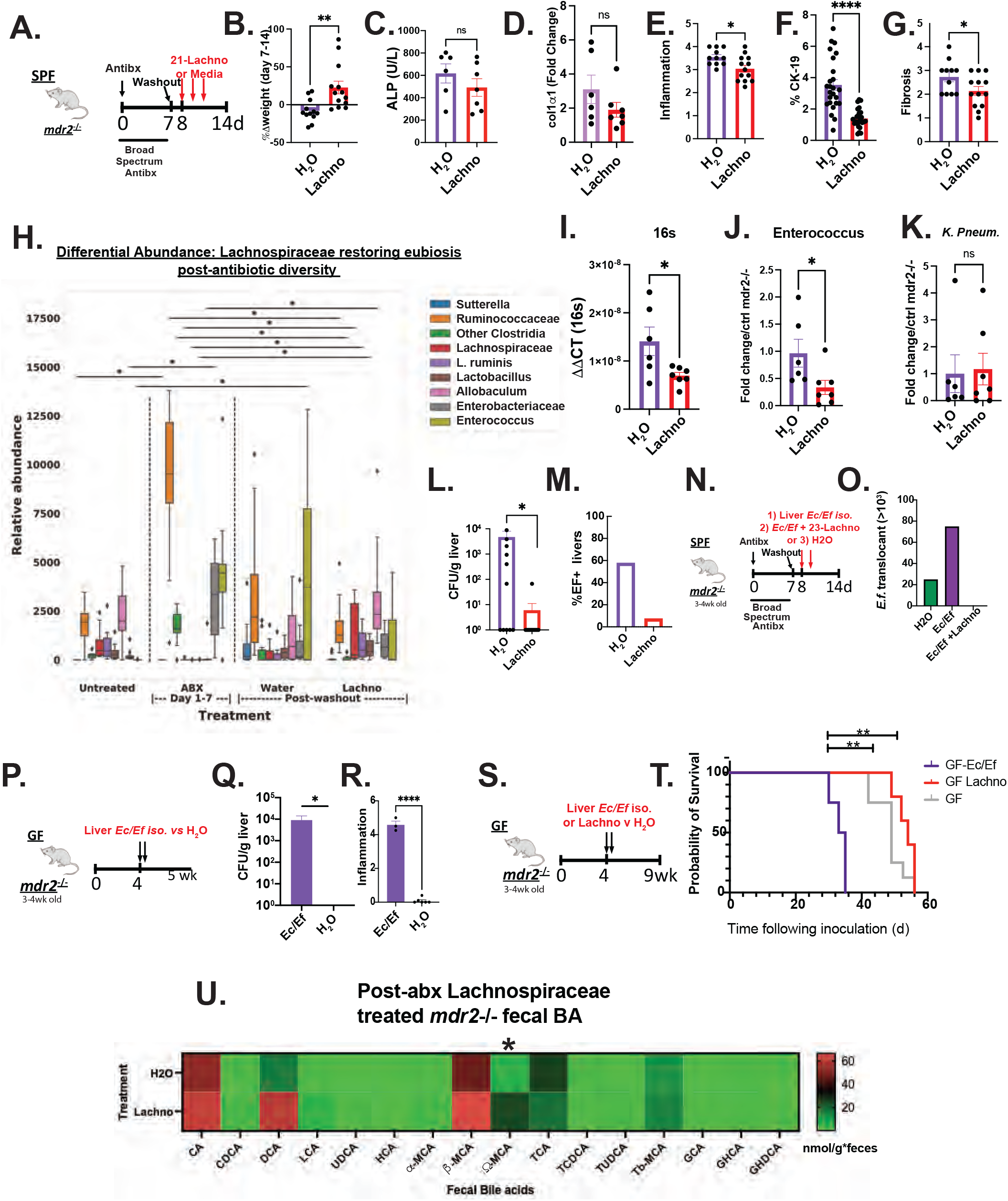
Lachnospiraceae (Lachno) phenotypic and metabolic effects on dysbiosis in *mdr2*^*-/-*^ mice. Using the 7d antibiotic pretreatment model, followed by a 1d washout, 21 Lachnospiraceae strains vs. H_2_0 were administered to SPF *mdr2*^*-/-*^ mice on d8, 10, and 12 (A). Following Lachno treatment change readouts included Δweight over 7 days (B), serum ALP (C), liver RNA col1α1 expression (D), blinded histologic inflammation, ductal reaction , hepatic fibrosis scoring (E-G). Differential abundance expression of 16s rRNA analysis (H) (N, *mdr2*^*-/-*^ untreated=24, Abx=36, H_2_O=11, Lachnospiraceae=13, pooled from 2 experiments. Presence of universal 16s, Enterococcus, and *Klebsiella pneumoniae* (K.P) fecal DNA by qPCR (I-K) (1 of 2 representative experiment, N=6 mice/group). Hepatic bacterial translocation (L), % of mice with *E. faecalis* liver translocation (M). Experimental design of treatment of 3-4wk old SPF *mdr2*^*-/-*^ mice treated with broad-spectrum antibiotic cocktail (vancomycin, neomycin, and metronidazole) for 7d followed by a 1d washout (N), then inoculated with *mdr2*^*-*/-^ resident Ec/Ef isolates, vs Ec/Ef + 23-Lachno combination or water only controls assessing translocated bacteria cultured from homogenized liver (O) (N=4 mice/group). Experimental design of treatment of 3-4wk old GF *mdr2*^*-/-*^ mice gavaged twice with pooled *E. faecalis* and *E. coli* hepatic isolates (Ec/Ef) 10^8^ CFU (P), measuring translocated liver bacteria (Q) and blinded liver inflammation score (R) of GF *mdr2*^*-/-*^ mice or water treated controls. Experiment design and Kaplan-Meyer survival curves of 10^8^Ec/Ef vs Lachnospiraceae (Lachno) or H2O controls (S-T) GF *mdr2*^*-/-*^ H2O, N=12; Ec/Ef, N= 5, Lachno, N=6). Heat map representation of cecal content bile acids (U) following Lachno vs H2O control *mdr2*^*-/-*^ accelerated antibiotic model. Group or pairwise comparisons performed by ANOVA or Student t-test, respectively. *P < .05, **P < .01, ***P <.001, ****P < .0001.

Longitudinal fecal microbial communities were altered 7 days after antibiotic treatment with expanded Ruminococcaceae, Enterobacteriaceae, and especially Enterococcus vs baseline untreated communities (Fig. 5H). Lachnospiraceae supplementation more closely resembled the pre-antibiotic state and particularly repressed Enterococcus expansion compared to water controls (Fig. 5H), confirmed by real-time PCR of fecal samples (Fig. 5I-J). Lachnospiraceae treatment did not decrease fecal *Klebsiella pneumoniae* ^2^ (Fig. 5K). Interestingly, Lachnospiraceae treatment reduced culturable hepatic translocated bacteria following antibiotic exposure (Fig. 5L). Sanger sequencing identified the translocating strains as *E. faecalis* and *E. coli*, with 60% of the translocants in Abx-treated SPF *mdr2*^*-/-*^ mice being *E. faecalis* (Fig. 4J); all hepatic *E. faecalis* isolates expressed the exotoxin cytolysin ^25^.

To more robustly assess hepatic changes directly caused by *E. faecalis* (*Ef*) and *E. coli* (*Ec*), 3-4wk old SPF-antibiotic pretreated or GF *mdr2*^*-/-*^ mice were colonized with selected *Ec/Ef* hepatic isolate strains or H2O and euthanized at 5wk of age (Fig. 5N,5P). We confirmed translocation of *Ec/Ef* liver isolates in both antibiotic pre-treated and gnotobiotic conditions (Fig 5O,5Q) relative to untreated controls in 5-week GF *mdr2*^*-/-*^ mice. Treatment with Lachnospiraceae prevented *Ec/Ef* liver translocation (Fig. 5O) similar to results seen in antibiotic-treated SPF mice (Fig. 5J). We assessed the lethality of these bacteria in gnotobiotic *mdr2*^*-/-*^ mice by orally inoculating *Ec/Ef* isolates with/without 23-Lachnospiraceae strains (Fig. 5S-T). The *Ec/Ef* dual-associated mice exhibited earlier mortality relative to H2O controls or Lachnospiraceae-colonized gnotobiotic *mdr2*^*-/-*^ mice (Fig. 5T).

### Lachnospiraceae repletion altered bile acid metabolism

Because antibiotics severely altered BA profiles, we assessed fecal bile metabolism following Lachnospiraceae administration to antibiotic treated *mdr2*^*-/-*^ mice. Lachnospiraceae significantly increased hydrophilic ω-MCA levels (Fig. 5U), a derivative of β-MCA conversion by 6-β−dehydroxylation. Lachnospiraceae treatment did not significantly alter total BAs (Fig. S2L), nor affect ileal FGF-15 or hepatic cyp7a-1 gene expression (S2M-N).

### SCFAs produced by Lachnospiraceae have hepatic anti-fibrogenic effects

We postulate these anti-inflammatory and anti-fibrotic effects are mediated by Lachnospiraceae metabolic products, including short-chain fatty acids (SCFAs), that are depleted by antibiotics (Fig. 6B). Vancomycin most effectively decreased SCFA (Fig. 6B), suggesting vancomycin-sensitive, resident bacteria, like Lachnospiraceae, are predominant SCFA producers. Oral supplementation of SCFAs (acetate, propionate, and butyrate) in vancomycin-treated *mdr2*^*-/-*^ mice attenuated histologic hepatic fibrosis and fibrosis gene expression (Fig. 6C-G), but had less impressive effects on inflammation. Lachnospiraceae strains with variable SCFA production were divided into 3 strains with relatively low (8, 9, 21-Lo) or high (52, 60, 70-Hi) SCFA production as determined by mass spectrometry of supernatants (Fig. 6H). We serially gavaged Hi or Lo SCFA-producing Lachnospiraceae for 6d to SPF *mdr2*^*-/-*^ mice after 7d antibiotic pretreatment, beginning after the washout period. (Fig. 6I). Hi SCFA-producing Lachnospiraceae attenuated histologic hepatic fibrosis (Fig. 6J) but not hepatic inflammation (Fig. 6K) compared with the low SCFA-producing group. These results provided a mechanism for protection by our Lachnospiraceae consortium.

**Figure 6:**
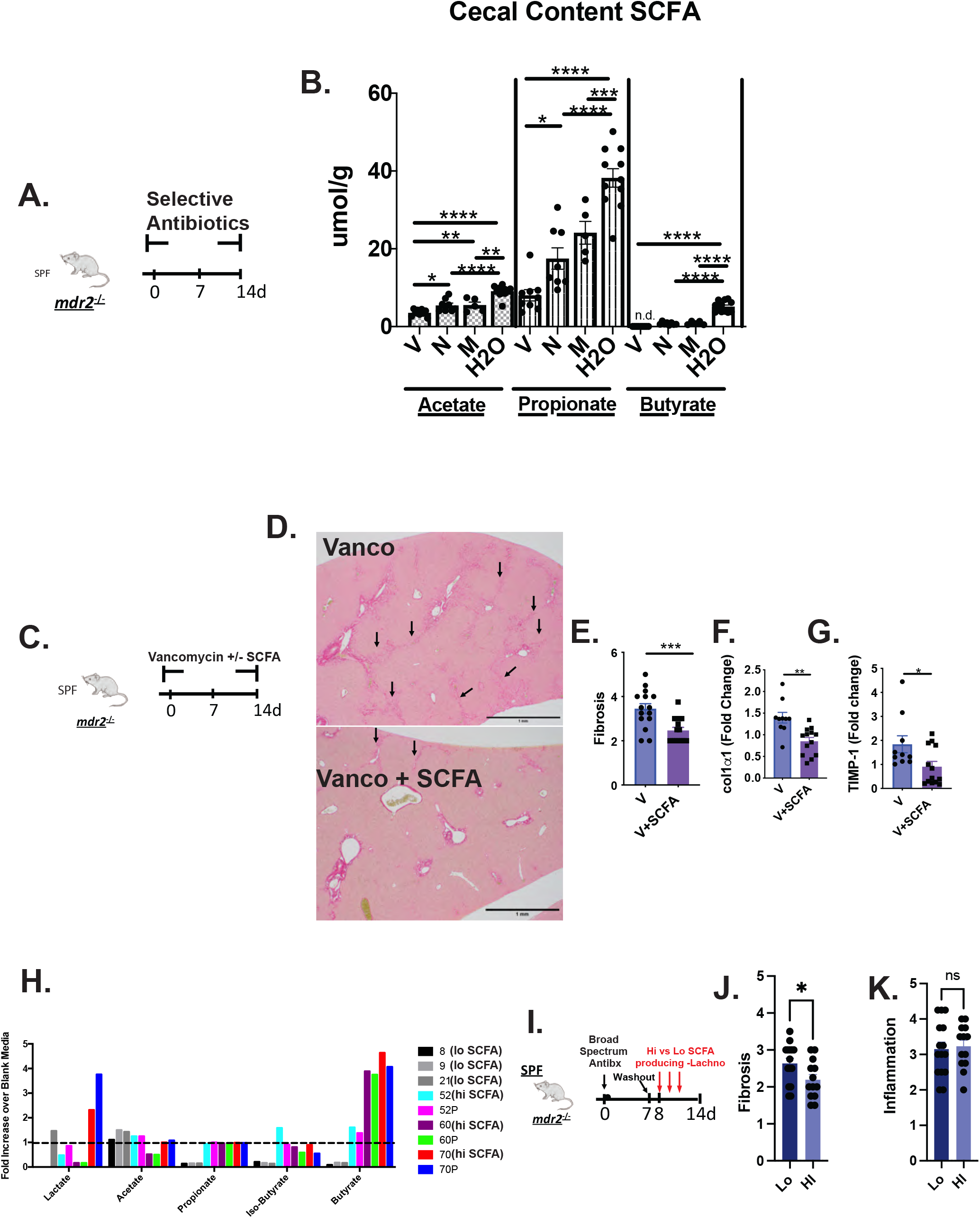
Effect of Short-Chain Fatty Acids (SCFA) on *mdr2*^*-/-*^ antibiotic model. Experimental design of 14d selective antibiotic treatment (A) with measurement of cecal content SCFA in SPF *mdr2*^*-/-*^ mice. Design of SPF *mdr2*^*-/-*^ mice treated with vancomycin (0.5 mg/ml in drinking water, N=10) with and without SCFA (67.5 mM acetate, 25.9 mM propionate, 40 mM butyrate and 3% sucrose, N=13) ad libitum, pooled from 2 experiments with representative Sirius Red photomicrographs of vancomycin vs. vancomycin + SCFA SPF *mdr2*^*-/-*^ mice, arrows highlight bridging fibrosis (D), blinded histologic fibrosis scoring (E) along with liver RNA col1α1 and *timp1* expression (F-G). Three relatively low and high SCFA-producing (Lo strain #s 8, 9, 21; Hi #s 52, 60, 70) Lachnospiraceae strains selected from 21 consortium strains were identified by mass spectrometry (H). Experimental design of accelerated antibiotic pretreatment of SPF *mdr2*^*-/-*^ mice for 7d and following a 1-day washout, administering Hi or Lo SCFA-producing Lachnospiraceae strains by gavage on days 8, 9 and 10, then followed for 6d (I) with measurement of histologic hepatic fibrosis (J) and inflammation (K) in Hi compared to Lo SCFA-producing strains. Group or pairwise comparisons performed by ANOVA or Student t-test, respectively. *P < .05, **P < .01, ***P <.001, ****P < .0001.

### *E. faecalis* and Lachnospiraceae are associated with divergent PSC clinical outcomes

Analysis of metagenomic data from multi-national repositories (Norway and Germany)^3^ of PSC fecal samples identified enrichment of *E. faecalis* and Enterobacteriaceae (Fig. 7A,B) and depletion of Lachnospiraceae (Fig. 7C) in PSC patients compared to healthy controls. A positive correlation existed between both *E. faecalis* (species) and Enterobacteriaceae (family) and Mayo risk score (n=51, rho 0.28 for both, p<0.05), with negative correlations between several Lachnospiraceae species and mayo risk score (rho ∼ -0.3, p<0.05) but only a trend at the family level (rho -0.2). *E. faecalis* is more prevalent in patients with PSC than controls (17% vs 5%) (Fig. 7D), which was amplified in patients who received antibiotics the last 6 months (Fig. 7D). Trends existed towards increased Enterobacteriaceae and reduced Lachnospiraceae in patients with antibiotics last 6 months. These clinical results mirrored observation in our *mdr2*^*-/-*^ mice providing clinical relevance to our murine results.

**Figure 7:**
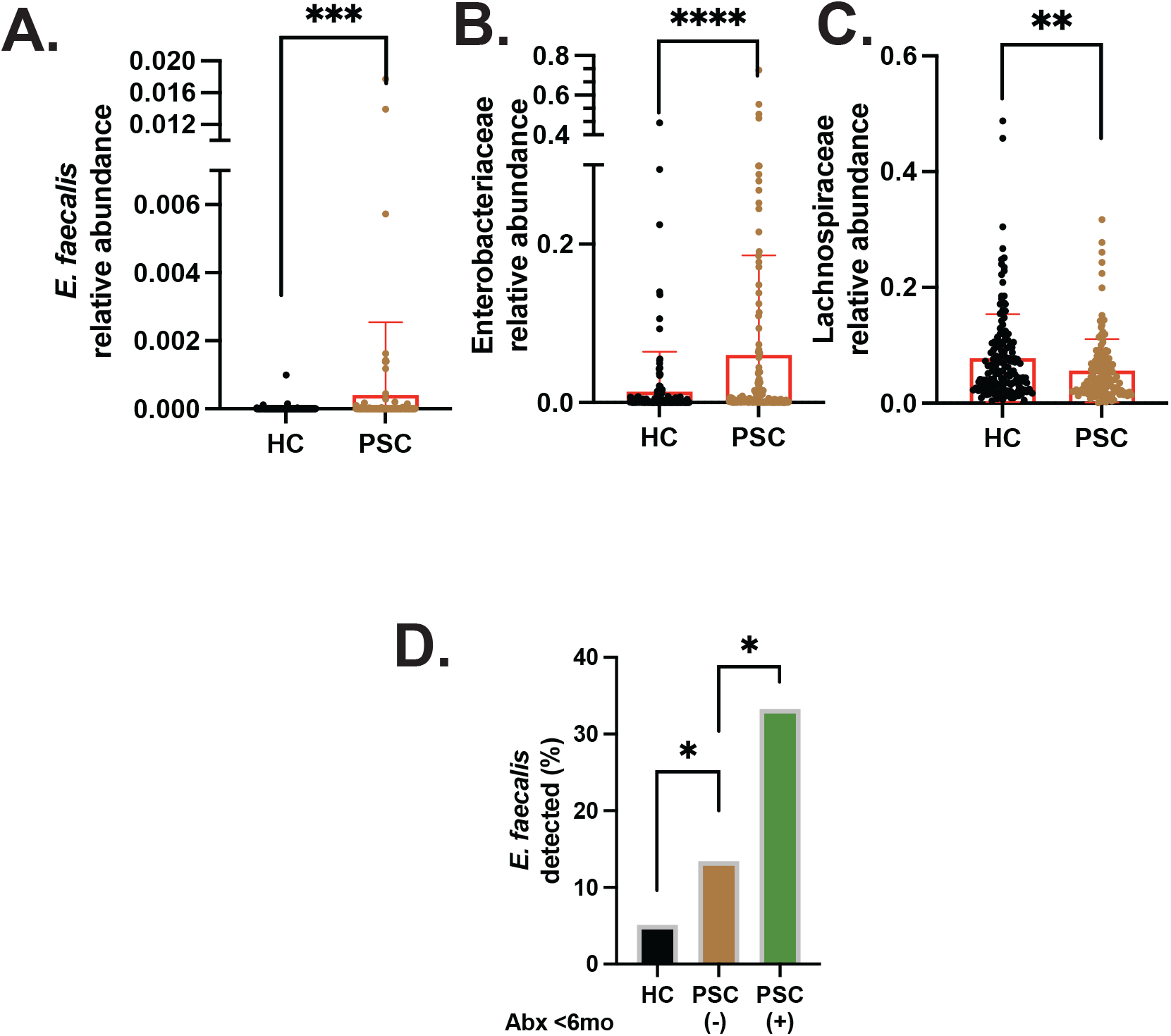
Selective fecal bacterial profile in PSC patients and effect of antibiotic exposure. Differential relative abundance of fecal *E. faecalis* (A), Enterobacteriaceae (B) and Lachnospiraceae (C) in heathy controls (HC, n-136) vs PSC (n = 158) patients. Prevalence of fecal *E. faecalis* detectable HC vs PSC patients exposed to antibiotics during the last 6 months. PSC no Abx (n=112) and with Abx (n=24) in last 6 months.

## Discussion

Our novel *in vivo* platform identified and tested functions of potential protective and detrimental resident bacterial groups to better understand and manipulate resident microbial drivers of protection and disease pathogenesis in a PSC model. We confirm the overall protection of resident microbiota in GF mice^13,12^ in a more aggressive genetic background of *mdr2*^*-/-*^ mice (C57BL/6)^26^. We observed early lethality and severe cholestasis in GF *mdr2*^*-/-*^ mice (median survival: 7.5wks) with TNF−α and Rorγt induction, robust periportal macrophage recruitment and accelerated liver inflammation and fibrosis. Early (<5wks) but not later (>6wks old) autologous SPF FMT rescued this lethal phenotype, indicating a strong protective role for SPF *mdr2*^*-/-*^ microbiota, but a brief therapeutic window. This brief window may result from rapidly advancing disease in GF *mdr2*^*-/-*^ mice after 5.5-6wks, and/or effective microbial-mediated protection requires longer microbial engraftment.

Broad-spectrum antibiotic treatment confirmed the overall protective role of microbiota. A novel aspect of our study is using selective antibiotics to identify protective and aggressive resident bacterial subsets. We narrowed the protective microbial community and bacterial metabolites to a vancomycin-sensitive population by 16S profiling and targeted metabolomics. Lachnospiraceae^9^, Clostridaeceae^22^ and BA and SCFA metabolic pathways were markedly depleted in vancomycin-treated SPF *mdr2*^*-/-*^ mice. Selective colonization of GF and antibiotic-depleted mice with a human Clostridaceae consortium with protective benefits in several colitis models^22^ did not confer protection in our model, but Lachnospiraceae attenuated both experimental hepatic inflammation and fibrosis in antibiotic pre-treated mice. Lachnospiraceae actively produce SCFA^27^, and SCFA supplemental administration decreased hepatic fibrogenesis in our model. Antifibrogenic effects of Lachnospiraceae strains that produce high amounts of SCFA, but not by low SCFA producing strains support that SCFA mediate in part Lachnospiraceae protective effects. Metagenomic analysis of an established PSC patient repository documented clinical relevance of our findings^4^, newly observing that Lachnospiraceae species are decreased in PSC patients^4,8^.

Parallel to identifying protective Lachnospiraceae, we determined that *E. coli* and *E. faecalis* potentiated experimental disease. Antibiotics increased Enterobacteriaceae and Enterococci fecal concentrations and induced *E. coli* and cytolysin^+^ *E. faecalis* hepatic translocation. Hepatic translocation of *E. faecalis* into the biliary epithelium and enteric Enterobacteriaceae are associated with detrimental inflammatory profile in PSC patients^28^,^5^. Translocation of *E. faecalis* producing cytolysin, a two-subunit exotoxin, strongly predicts adverse outcomes in clinical studies and murine modeling of alcohol-associated hepatitis^25^. Importantly, Lachnospiraceae repletion of *mdr2*^*-/-*^ mice following antibiotic pretreatment inhibited *E. faecalis*/Enterobacteriaceae colonization and hepatic translocation. Accelerated mortality following selective colonization of GF *mdr2*^*-/-*^ mice confirmed direct pathogenicity of the translocating murine *E. faecalis/E. coli* strains. We documented increased *E. coli* and *E. faecalis* in a subset of PSC patients and their first published association with poor clinical outcomes.

Assessing cecal BA profiles by multiple parameters, including total BA, 7-α-dehydroxylation activity, taurine-conjugated α+β muricholic acids and FXR signaling following selective antibiotic treatment in SPF *mdr2*^*-/-*^ mice demonstrated a vancomycin-specific increase in unconjugated/conjugated BA, a marker of BSH activity. A recent 16s rRNA analysis of biliary fluid demonstrated an increased *E. faecalis* in PSC patients associated with increased unconjugated/conjugated BA relative to controls, suggesting increased BSH activity^3^. Unconjugated (less hydroxylated) hydrophobic BA can injure hepatocytes^29^ and cholangiocytes^30^, providing a mechanism of accelerated disease with Abx. Lastly, increased hydrophilic BA UDCA in neomycin-treated *mdr2*^*-/-*^ may partially explain our observed lack of potentiated hepatobiliary pathology, due to the protective effect of UCDA in human PC and experimental models^14 31 32^. These results support our model’s translational potential for identifying potential biomarkers and therapeutic metabolites to guide clinical studies.

Small clinical trials of different antibiotics, including vancomycin, in PSC patients report short-term efficacy by biochemical outcomes^33^. However, up to 1/3-1/2 of patients are non-responders^33 34^. Possible explanations for non-responding subsets include: 1) resistant aggressive bacteria, 2) narrow-spectrum antibiotics not targeting an individual’s dominant pathobionts, 3) non-specific depletion of protective commensal species and 4) non-microbial mediated disease. A dearth of pre- and post-intervention microbial profiling failed to elucidate these trials’ mechanisms of success vs. failure. Our human and murine data demonstrate that the deleterious dysbiotic effects of antibiotics depend on several factors: 1) microbial spectrum and 2) host baseline resident bacterial constituency (presence or absence of pathogenic/protective bacteria). Future studies need to identify subgroups of PSC patients who may optimally benefit from antibiotic therapy along with more definitive understanding of microbial drivers and inhibitors of clinical inflammation and fibrosis, perhaps guided by observations in experimental models.

This study has several limitations. Early lethality of our GF and broad-spectrum antibiotic-depletion *mdr2*^*-/-*^ mice precluded longitudinal studies to assess delayed effects of Lachnospiraceae or SCFA on chronic disease. Secondly, we acknowledge the ongoing controversy regarding suitability of murine AP to monitor cholestasis^35,36^, and therefore utilized additional markers of cholestatic injury, TB and bile duct reactivity, to confirm our findings. Finally, because GF *mdr2*^*-/-*^ mice, particularly females, expire before becoming fertile, our heterozygous *mdr*^*+/-*^ breeding strategy with small litter sizes delayed experiment and limited more in-depth mechanistic studies, but permitted comparing littermate controls.

Our mechanistic studies have direct translational implications for future personalized bacterial manipulation in PSC patients. For example, our results might inform selection of FMT donors and recipients and guide strategies to selectively enrich Lachnospiraceae and deplete of *E. faecalis* and *E. coli* in clinical trials. Additionally, our axenic model can explore hepatobiliary, microbial, and metabolomic impacts of humanized fecal transfer and selective WT and genetically engineered bacterial strain colonization to dissect mechanisms of microbial protection and pathogenesis. Future directions include better understanding the influence of macrophage phenotypes^37^, microbial BSH-mediated effects, and perhaps other putative protective bacteria on hepatobiliary inflammation and fibrosis. In conclusion, our axenic and antibiotic *mdr2*^*-/-*^ PSC murine models identified complex interactions between novel hepatoprotective microbes balanced against pathobionts, replicated in PSC patient cohorts. Resident microbes exert both positive and negative influences partly through anti-microbial activities and altering bacterial metabolites.

## Grant support

NIH grants P01DK094779, P40OD10995, P30DK034987 (to RBS), T32DK07737 and T322A1007273 (to MA), The Crohn’s and Colitis Foundation #2434 (to RBS); CA232109, DK094779, AI067798 (to JT); K01DK119582 (to JW); P30 DK120515 (to BS), 1I01BX003031; DK108959 and DK119421 (to HF). JRH is funded by the European Research Council, grant no. 802544. CS is supported by the DFG, CRU306, and by the Helmut and Hannelore Greve Foundation and grant support from BiomX and Galapagos.

## Abbreviations

Abx: broad-spectrum antibiotics
ALP: alkaline phosphatase
ALT: alanine aminotransferase
BA: bile acids
BSH: bile salt hydrolase
CA: Cholic Acid
CDCA: chenodeoxycholic acid
*col1a1*: collagen 1a1
cyto: cytolysin
DCA: Deoxycholic Acid
FMT: fecal microbiota transplant
GF: germ-free
HYP: hepatic hydroxyproline
Lachno: Lachnospiraceae
LC: Lithocholic Acid
Lcn-2: lipocalin-2
M: metronidazole
MCP: monocyte chemoattractant protein
*mdr2*^*-/-*^: multi-drug resistant 2 gene knockout
MCA: muricholic acid
N: neomycin
OTU: operational taxonomic unit
PSC: Primary Sclerosing Cholangitis
SCFA: short-chain fatty acids
SPF: specific pathogen-free
TIMP-1: tissue inhibitor of metalloproteinase-1
TB: Total bilirubin
V: vancomycin
WT: wild-type

## Disclosures

RVS has consulted for and received grant support from Takeda, Janssen, Second Genome, Vedanta, BiomX, Biomica, SERES and Artizan; B.S. has been consulting for Ferring Research Institute, HOST Therabiomics, Intercept Pharmaceuticals, Mabwell Therapeutics, Patara Pharmaceuticals and Takeda. B.S.’s institution UC San Diego has received grant support from Axial Biotherapeutics, BiomX, CymaBay Therapeutics, NGM Biopharmaceuticals, Prodigy Biotech and Synlogic Operating Company. B.S. is founder of Nterica Bio.

## Writing Assistance

none

## Authors’ Contributions

MA and RBS conceptualized the study. MA, BN, AV, MF contributed to the collection of samples. JT, JPYT grew and contributed Lachnospiraceae strains. KL and YL were responsible for metabolomics. JW performed the post-sequencing data processing. SAM performed blinded histological scoring and MA executed histologic analysis. MK, LT, CS, CB, AF, JH collected and analyzed human fecal metagenomic data. BS provided technical advice, reviewed data and edited the results. MA, JW, and RBS planned the analysis. MA, JW, and RBS performed data curation and formal analysis. MA and RBS interpreted data. RBS supervised the study, and MA was responsible for project administration. MA and RBS wrote the original draft. MA, JW, RBS, and KL contributed to the writing of the manuscript. All authors read, critically revised for important intellectual content, and approved the final manuscript. RBS and MA acquired funding.

## Acknowledgments

Bo Liu and Akihiko Oka, MD provided technical assistance; Josh Frost, UNC Gnotobiotic Facility Manager for gnotobiotic research support; Fengling Li, MD, Ph.D., and Allison Rogala, DVM for germ-free derivation of mdr2^-/-^ mice; Lisa Holt for laboratory management and Cary Cotton for statistical support. Special appreciation to Vincent Young, University of Michigan, for providing the original Lachnospiraceae stains.

## Notes

### Competing Interest Statement

The authors have declared no competing interest.

