## Supplementary figures and images for "Protective and aggressive bacterial subsets and metabolites modify hepatobiliary inflammation and fibrosis in PSC"

### Supplemental figures

#### Supplemental Figure 1

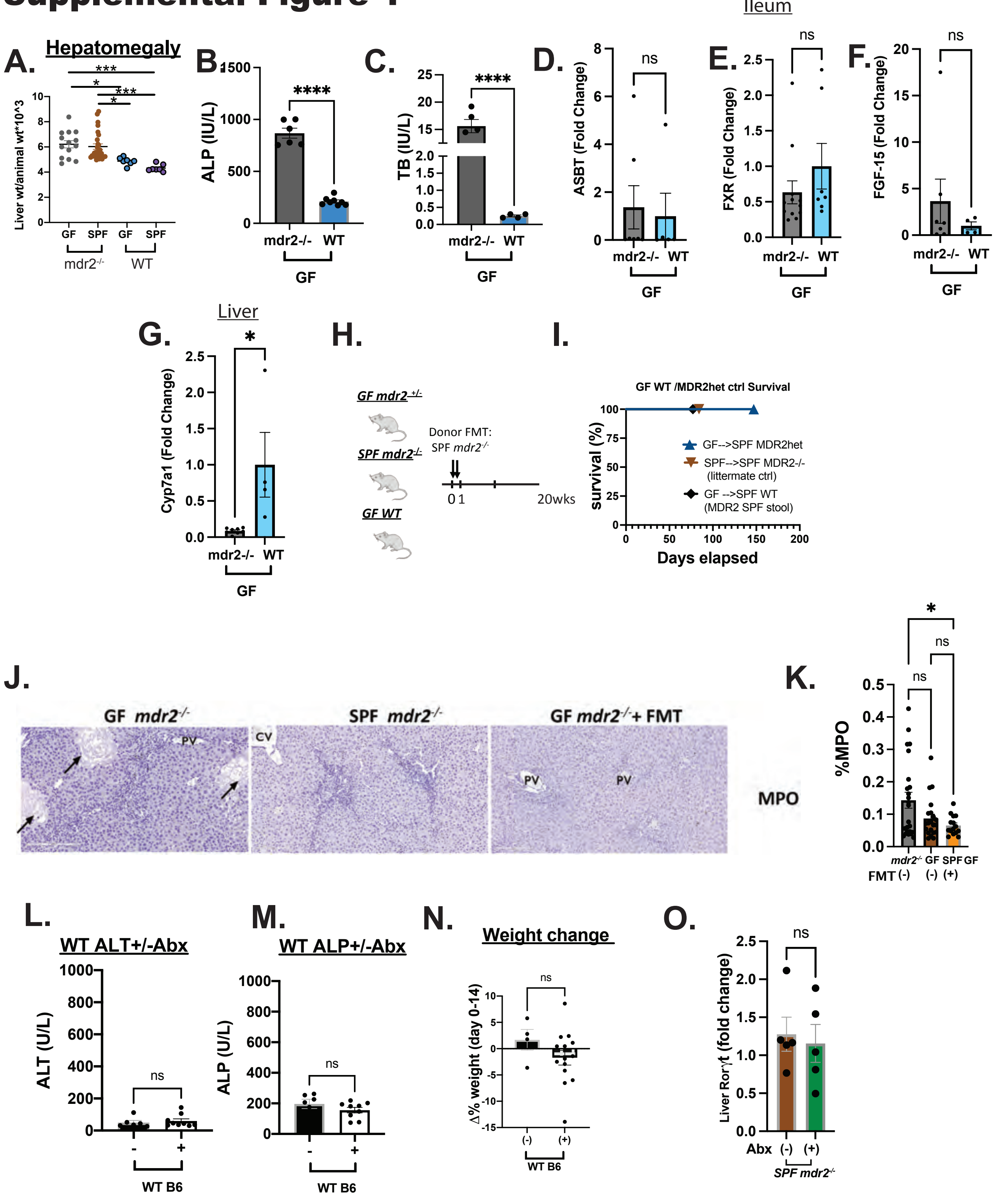

#### Supplemental Figure 2

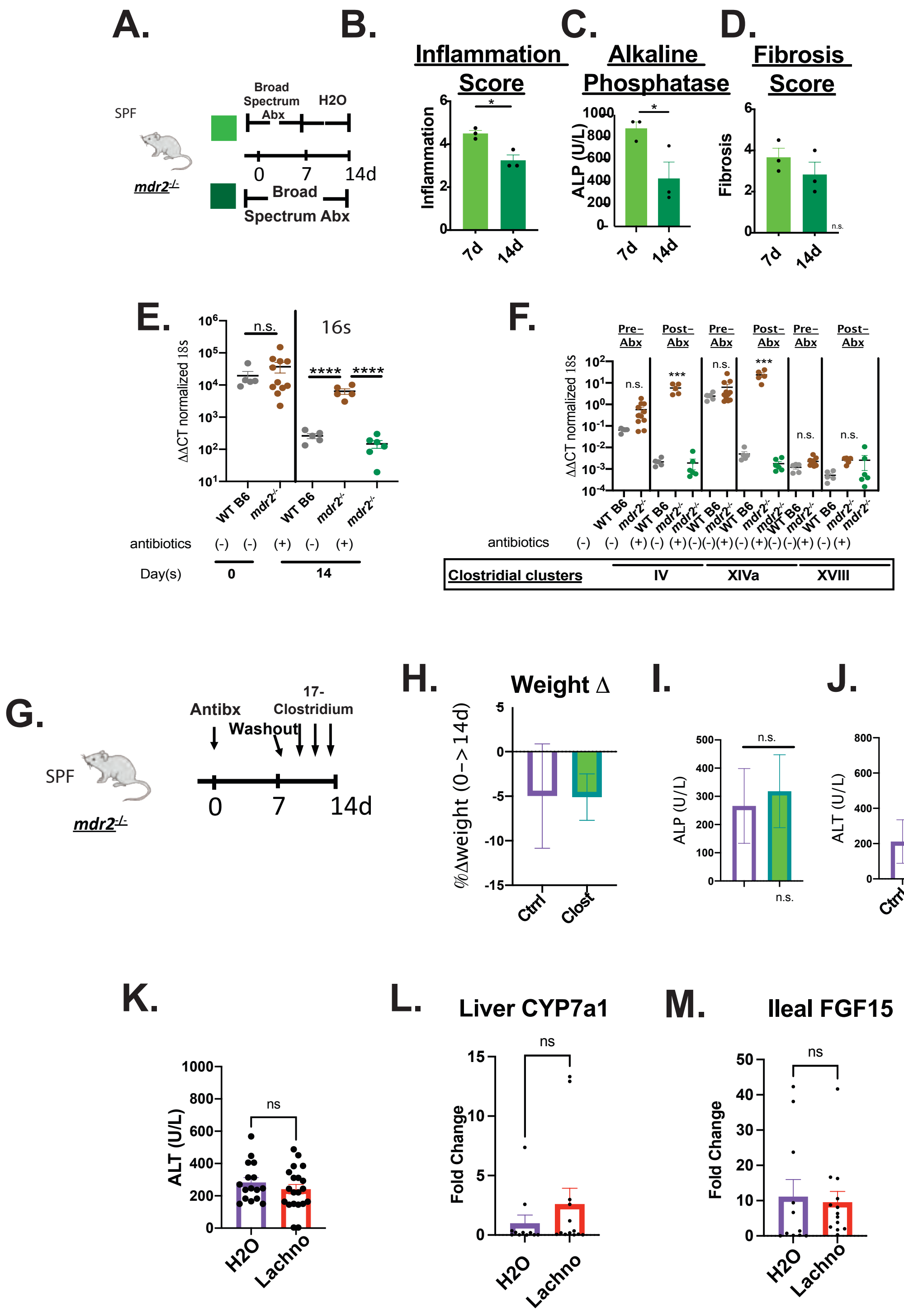
