## supplemental figure legends for "Protective and aggressive bacterial subsets and metabolites modify hepatobiliary inflammation and fibrosis in PSC"

**Supplemental Figure 1:** **Characteristics of GF/ SPF *mdr2^-/-^*and WT mice with and without broad-spectrum antibiotics.** Differences in liver weight amongst 6-8wk old GF and SPF wild-type and *mdr2^-/-^* mice (A). Pooled serum ALP (B) and TB(C), ileal expression of apical sodium dependent bile acid transporter (ASBT), farsenoid-X receptor (D, FXR), fibroblast growth factor 15 (E, FGF-15), along with liver cyp7a1 (G) in GF *mdr2^-/-^* vs WT mice. Survival in untreated SPF WT and *mdr2^-/-^* mice (WT= 13, *mdr2^-/-^* 23) (H-I). Representative photomicrographs of 6-8wk old GF, SPF, or post-FMT GF *mdr2^-/-^* murine liver stained by myeloperoxidase (MPO) (J, 100X) along with composite automated scoring of % MPO positive cells. Serum ALT (L), ALP(M) and 14d change in weight (N) following oral gavage of broad-spectrum antibiotics (vancomycin, neomycin and metronidazole) ad libitum compared to water controls. Hepatic expression of rorgt following broad spectrum abx for 14days (O). Results are expressed as means +/- SEM. Survival data analyzed by Log-rank (Mantel-Cox) test, group or pairwise comparisons performed by ANOVA or Student t-test, respectively. PCoA of the beta were analyzed by permanova analysis. P-value*P < .05, **P < .01, ***P <.001, ****P < .0001.

**Supplemental Figure 2:** **Efficacy of putative hepatoprotective resident bacteria in *mdr2^-/-^* mice.** Experimental design of accelerated antibiotic treatment model: SPF *mdr2^-/-^* mice were treated with broad-spectrum antibiotics (vancomycin, metronidazole and neomycin) for 7d vs. 14d (A) resulting in aggressive liver inflammation and injury: the 7d antibiotic regimen had increased histologic inflammation (B) and serum alkaline phosphatase (C), but not liver fibrosis (D) (7d, N=3, 14d, N=3, 2 experiments). Qpcr amplified results of fecal samples from *mdr2^-/-^* mice with and without broad-spectrum antibiotics (vancomycin, neomycin, and metronidazole) using the 16s (E) and Clostridial cluster primers (cluster IV, XIVa, and XVIII) (F) normalized to 18s. Experimental design of treatment of 3-4wk old SPF *mdr2^-/-^* mice treated with 7 day treatment of broad antibiotic cocktail followed by a 1d washout period, then the inoculation of 17-strain of Clostridium (G) and assessing 14 weight change (H), ALP & ALT (I-J). (n=4 mice in each treatment and control groups). Following Lachnospiraceae treatment of *mdr2^-/-^* mice outlined in Fig4E (lachno, N=13, H2O ctrl, N=12), we assessed pooled data from 2 separate experiments of ALT (K). qPCR assessment of orphan receptor FXR pathway by look at liver cyp7a1 (L), ileal FGF15 (M) in *mdr2^-/-^* exposed to Lachnospiraceae. Group or pairwise comparisons performed by ANOVA or Student t-test, respectively. *P < .05, **P < .01, ***P <.001, ****P < .0001.
